# Real Time Ultrasound Molecular Imaging of Prostate Cancer with PSMA-targeted Nanobubbles

**DOI:** 10.1101/634444

**Authors:** Reshani Perera, Al de Leon, Xinning Wang, Yu Wang, Gopal Ramamurthy, Pubudu Peiris, Eric Abenojar, James P. Basilion, Agata A. Exner

## Abstract

Contrast-enhanced ultrasound with microbubbles has shown promise in detection of prostate cancer (PCa), but sensitivity and specificity of detection remain challenging. Targeted nanoscale contrast agents with improved capability to accumulate in tumors may result in prolonged signal enhancement and improved detection of PCa with ultrasound. Here we report on a new nanobubble contrast agent that specifically targets prostate specific membrane antigen (PSMA) overexpressed in most prostate tumors. The PSMA-targeted bubbles (PSMA-NB) were utilized to simultaneously image dual flank PCa tumors (PSMA-positive PC3pip and PSMA-negative PC3flu) to examine whether the biomarker can be successfully detected and imaged using this probe in a mouse model. Results demonstrate that active targeting of NBs to PSMA rapidly and selectively enhances tumor accumulation and is critical for tumor retention of the contrast agent. Importantly, these processes could be visualized and quantified, in real time, with standard clinical ultrasound. Such demonstration of the immense yet underutilized potential of ultrasound in the area of molecular imaging can open the door to future opportunities for improving sensitivity and specificity of cancer detection using parametric NB-enhanced ultrasound imaging.

---

Despite significant efforts, prostate cancer (PCa) is still the second most common leading cause of cancer-related deaths worldwide, with 180,000 new cases diagnosed in the USA in 2018^1–2^. Accurate diagnosis of PCa is a crucial step necessary for informing the clinical management of the disease, yet conventional options leave much space for improvement. Currently, men with an abnormal digital rectal exam and/or increased levels of prostate serum antigen (PSA) are considered at high risk for cancer and are referred for a prostate biopsy to assess if PCa is present. The standard PCa biopsy procedure uses transrectal ultrasound (US) guidance to determine the prostate gland orientation, but the delineation of tumors within the prostate using US is unclear. Accordingly, biopsies are performed in a systematic manner by selecting 6-12 or more area from the peripheral zone of the prostate. These cores represent only 1% of prostate tissue and are a gross under sampling of prostate gland tissue, and biopsies performed using this conventional procedure result in significant false negatives of up to 50%^3–5^. Concern over the lack of pathological data in the face of other positive clinical risk factors results in almost 50% of patients undergoing second, if not third and fourth, prostate biopsies leading to increased costs and risk associated with unnecessary procedures. If the already-on board US technology can be used to more reliably identify the location of prostate cancer within the prostate gland, these outcomes stand to be significantly improved.

Contrast-enhanced ultrasound has been investigated as one option for improved PCa detection^6–7^. In order to increase the PCa detection rate while limiting the number of biopsy procedures, significant effort has been focused on formulation of lipid and/or protein-stabilized gas filled contrast agents to improve the US imaging capability of cancer within the prostate^8–10^. Most of these efforts have utilized micron-sized contrast agents or microbubbles (MBs)^9, 11–12^, which are already clinically utilized for other applications, but these have lacked specificity and sensitivity over conventional methods. One option to improve these parameters is molecular targeting of microbubbles to vascular biomarkers. One example of this approach currently in clinical trials is BR55 (Bracco, Geneva, Switzerland) ^13–15^, which is targeted to vascular endothelial growth factor receptor 2 (VEGFR2). BR55 was examined recently in a phase 0 study for its ability to detect PCa^16^. The reported detection of malignant lesions with BR55 was 68%. {Smeenge, 2017 #42}. Two factors may confound the use of MBs for this application. The first is their large footprint, which confines MBs to the blood stream, and makes consistent targeting and retention at vascular markers. It also makes most biomarkers for PCa and other cancer inaccessible, since most lie beyond the vasculature in the tumor parenchyma. Second, MBs have a short lifespan (typically < 10 min) in the circulation, making their utility limited during targeting and during the entire biopsy procedure.

These unmet needs have led to increased interest in nano-sized US contrast agents that can penetrate the tumor parenchyma, bind to cancers, and exhibit nonlinear contrast behavior similar to MBs^18–23^. Our group has recently developed an ultrastable nanobubble contrast agent (NB) that contains perfluoropropane inside a propylene glycol and glycerol-enhanced lipid shell, which demonstrated unique physicochemical properties and an extended life span *in vivo^24^*. These NBs should have the ability to pass though hyper-permeable vasculature in tumors^25–27^. Furthermore, active targeting to a biomarker overexpressed on prostate cancer should enhance the retention of administered NBs at the tumor site, thus facilitating enhanced molecular contrast imaging in the tumor. A well-known biomarker for PCa is the prostate specific membrane antigen (PSMA)^28–30^. PSMA is a type II integral membrane protein that is expressed at lower levels in the healthy prostate and other organs such as the kidney, liver, and brain, but significantly at higher levels in PCa^31–32^. The level of PSMA has a positive correlation with the pathological phase of the tumor stage^33–35^ and is also considered to be the most important protein target in diagnostic specific immunolocalization imaging and immune-directed therapy^31, 33, 36^. Many ligands specific to PSMA are available, including monoclonal and engineered antibodies, small sized molecules, nanobodies, and aptamers^9–10^

In the current study, we demonstrate the use of standard nonlinear contrast-enhanced US for real-time molecular imaging of PCa by targeting stable NBs to PSMA via the PSMA-1ligand, which has been previously validated as a robust marker for PCa^37–38^ (Fig. 1a). Our data demonstrate significant, sustained differences between the kinetics of PSMA-NB, NB and MB in PCa models. Specifically, the PSMA-NBs were rapidly taken up by the PSMA-expressing tumors and were selectively retained within the tumor parenchyma, significantly extending duration the signal available for US visualization compared to untargeted NBs and the commercially available MB, Lumason®. No differences were seen in the PSMA-negative tumors or kidneys between NBs and PSMA-NBs, suggesting that the observed retention is due to specific molecular targeting to PSMA in the tumor itself. Notably, since clinical US was consistently capable of distinguishing these kinetic differences in small tumors, the work lays the foundation for future work utilizing the technique in real-time US biopsy guidance.

**Figure 1.**
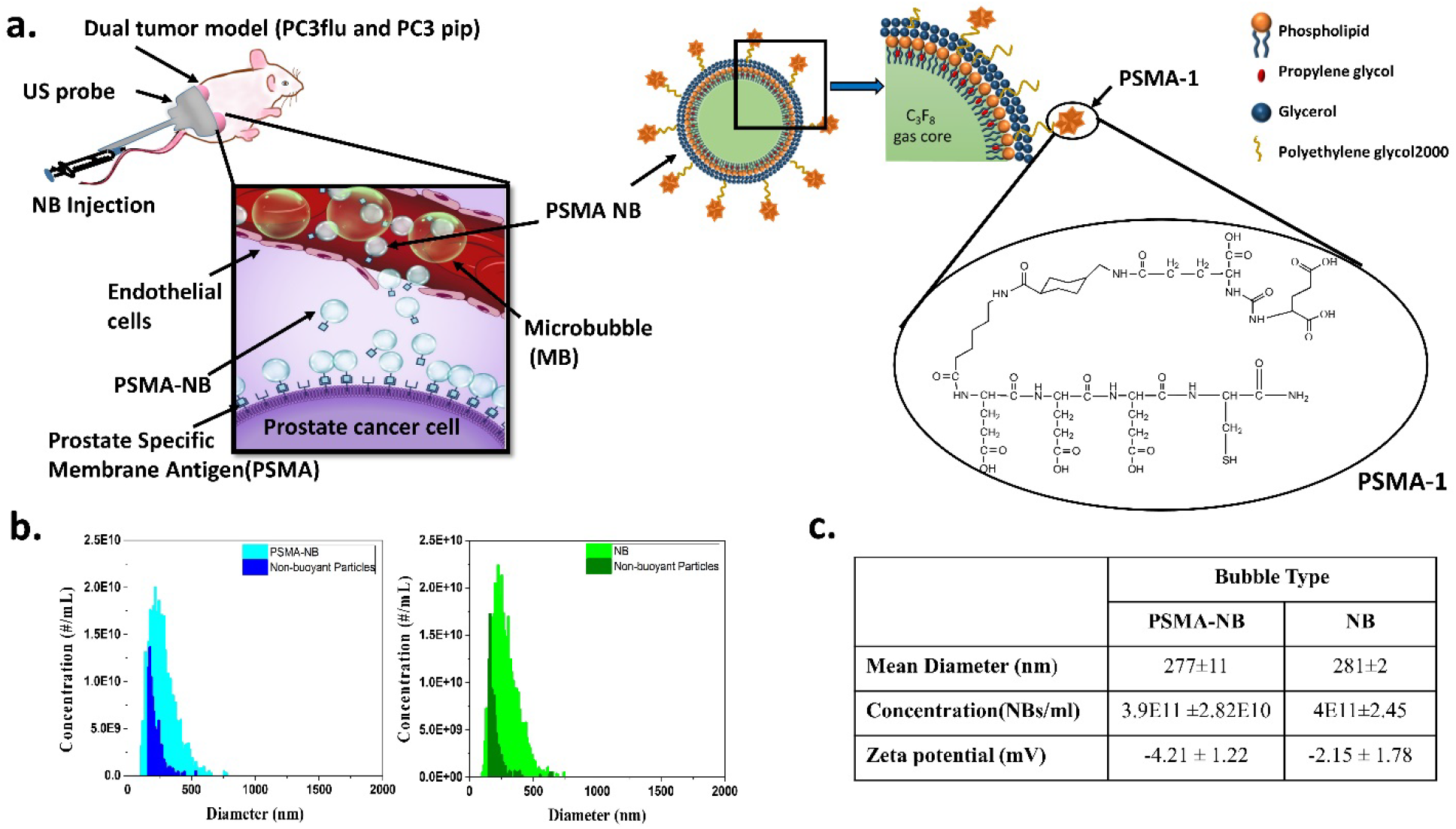
Design and characterization of PSMA-functionalized NBs for *in vivo* US molecular imaging. (a) Illustration of the experimental setup and schematic illustrating the delivery of PSMA-NB to the tumor parenchyma via their leaky vasculature after intravenous injection. DSPE-PEG-PSMA-1 is incorporated into the lipid shell of NB for targeting PCa cells that overexpress PSMA. (b) Size distribution and the concentration of NB and PSMA-NB acquired via resonant mass measurement. (c) The size, concentration, and the Zeta potential of bubbles. Conjugation of PSMA to the NBs has a minimal effect on the NB properties. Image in panel (a) based on art created by Erika Woodrum.

## Results

### Validation and Characterization of PSMA Targeted NB

The conjugation of PSMA-1 to lipids was confirmed by the HPLC and the MALDI-TOF-MS techniques (**Supplementary Fig. 1.a, b**). HPLC data showed that the PSMA-1 peak disappeared after conjugating PSMA-1 with DSPE lipid. MALDI-TOF-MS results further confirmed the conjugation of PSMA-1 to the DSPE lipid. The preparation and characterization of base ultrastable NBs has been reported elsewhere^24^. Briefly, the NB hydrodynamic diameter and concentration were characterized using resonant mass measurement (RMM) capable of detecting both buoyant (bubbles) and non-buoyant particles (liposomes, micelles, lipid debris, etc.). The RMM technology operates by measuring the change in the frequency of oscillation of particles that flow through an oscillating cantilever^24, 39–41^. The size and the concentration of PSMA-NB were 277±11 nm and 3.9E11 ±2.82E10 NBs/ml respectively. The mean size and the concentration did not change significantly after DSPE-PSMA was incorporated into the NBs (Fig.1b and Fig. 1c). Apart from gas filled bubbles, we also detected non-buoyant particles in both bubble solutions, which are invisible under US, but may contribute to the bubble overall stability^42^ (Fig. 1b). The slightly negative values of zeta potential crucial to the stability of both targeted and untargeted NB (Fig. 1c) were also confirmed using DLS measurement.

### *In vitro* cellular uptake studies

To optimize the amount of PSMA-1 ligand on the surface of NBs, bubbles with different amounts of PSMA-1 ligand were prepared, and labeled with Rhodamine B. The highest fluorescence was found in NBs labeled with reactions that contained 25 μg (35×10^3^ PSMA molecules per NB) of PSMA-1, followed by NBs with 50 μg (70×10^3^ PSMA molecules per NB) of PSMA-1, then decreased significantly in NBs with 100 μg (14×10^4^ PSMA molecules per NB) of PSMA-1 (**Supplementary Fig. 2a, b**). Our results concurred with other reports that best targeting was observed in nanoparticles with intermediate number of ligands per nanoparticle^43^. After the optimal ligand density was established, we then compared the cellular uptake of Rhodamine-PSMA-NB in PSMA-positive (PC3pip) cells and PSMA-negative (PC3flu) cells as shown in Fig. 2a. When cells were exposed to Rhodamine-NB, similar fluorescence intensity was observed in PC3pip and PC3flu cells. When PC3flu cells were exposed to Rhodamine-PSMA-NB, the fluorescent intensity was not significantly different compared to cells exposed to Rhodamine-NB (Fig. 2b). In contrast, the signal increased dramatically in PC3pip cells (~10 fold increased), which express PSMA, when treated with Rhodamine-PSMA-NB, indicating that PSMA-NBs selectively bind to PSMA-expressing cells.

**Figure 2.**
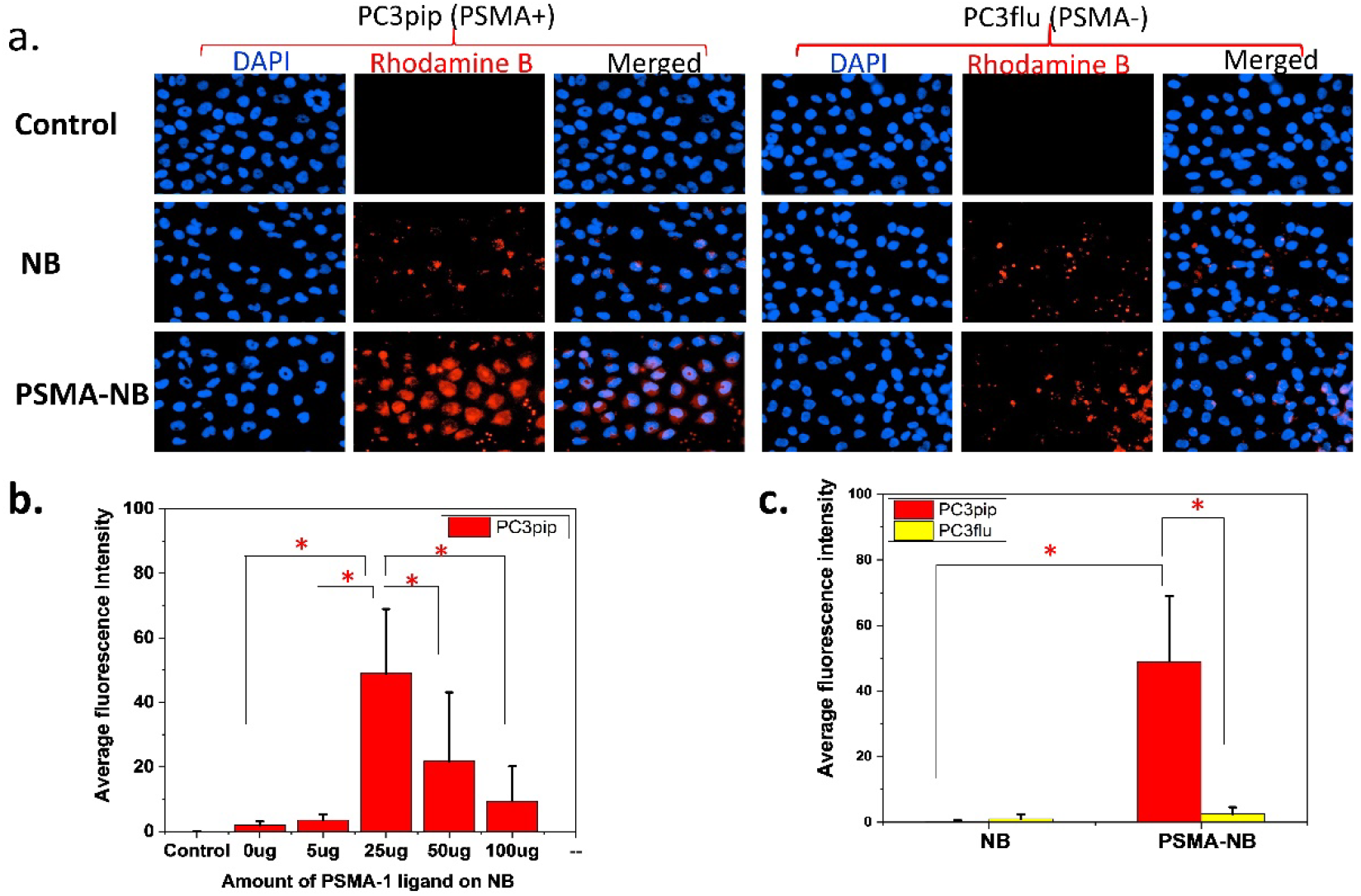
*In vitro* cellular uptake experiments reveal PSMA-NB selectively bind to the PSMA -positive PC3pip cells. (a) PSMA-positive PC3pip cells and PSMA-negative PC3flu cells on coverslips were incubated with no NB (control), Rhodamine-NB or Rhodamine-PSMA-NB for 1hr. Nuclei were stained using DAPI (blue) and uptake of Rhodamine tagged NB and PSMA-NB (red) was assessed by fluorescence microscopy. Images were taken at 40X. Representative images are shown from three independent experiments. (b) Quantification of fluorescence signal for bubbles with and without 25 µg of PSMA confirms significant increase in cell specificity with PSMA-NB showing >10 fold increase. n=3, error bars represent mean ± s.d., * P < 0.001.

### *In vivo* ultrasound imaging

*In vivo* experiments examining the kinetics of targeted PSMA-NB and NB were performed using clinical nonlinear contrast enhanced ultrasound and the average results from 7 mice are reported (Fig. 3.). Five quantitative parameters related to perfusion and bubble dispersion were extracted from the acquired time-intensity-curves (TIC). These include peak intensity, time to peak, the duration of contrast enhancement, and area under the curve. The parameters were compared between PSMA-NB and NB in the PSMA-positive PC3pip and PSMA-negative PC3flu tumors. Additional comparison was made with the commercially available microbubble, Lumason®.

**Figure 3.**
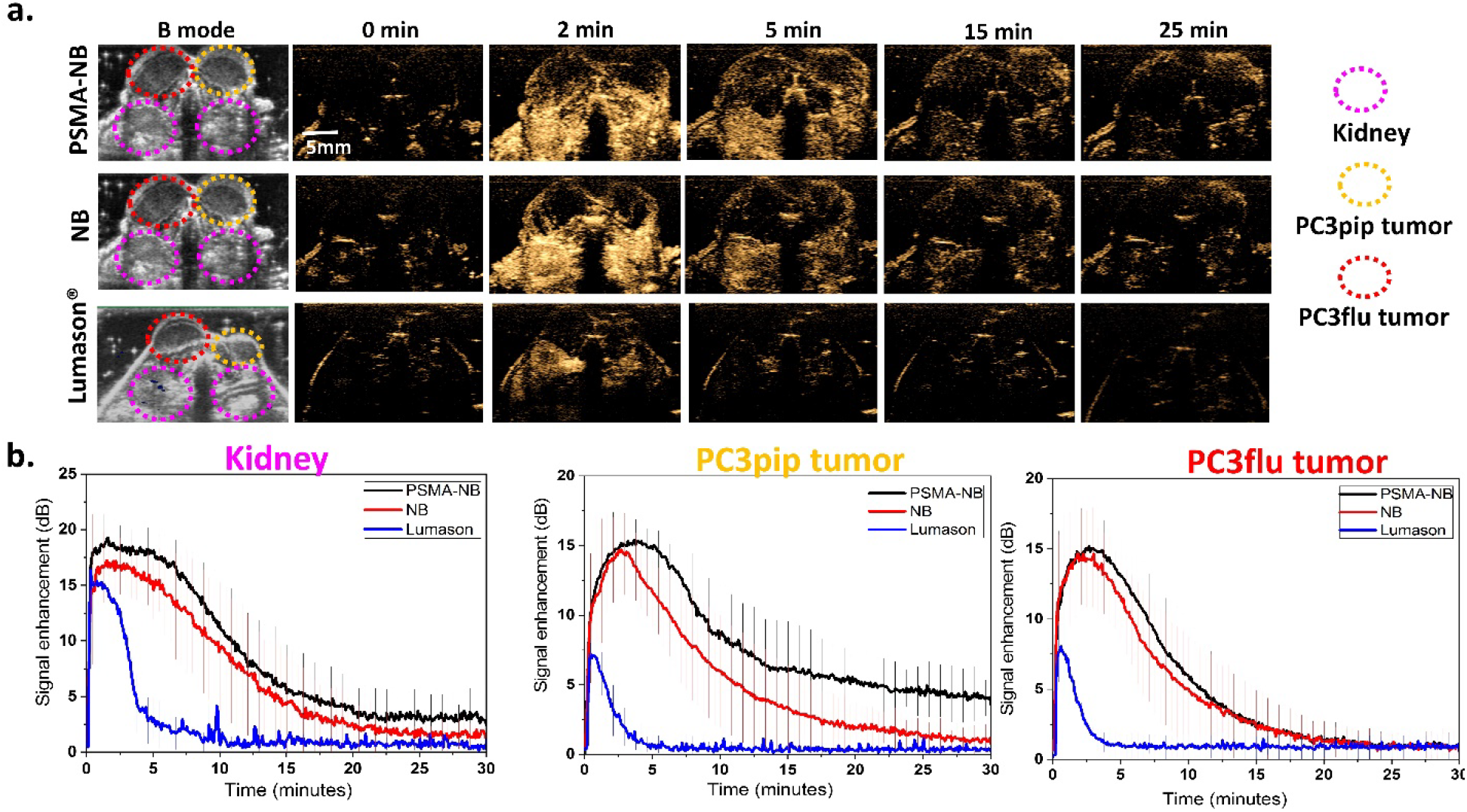
PSMA-NB enabled imaging of prolonged enhanced US signal in PSMA-positive PC3pip tumors. (a) Representative US imaging results of the targeted PSMA-NB, NB and Lumason® in PC3pip tumor, PC3flu tumor and kidneys. Bubbles were administered via tail vein and both PC3pip and PC3flu tumors and kidneys were imaged at 12 MHz, 245 kPa pressure, and 0.2 frames per second for 30 min. Left two columns show the B-mode and contrast harmonic imaging (CHI) mode images of tumors and kidneys before injection. The series of images shows the CHI images at different time point after bubble administration. At the peak intensity, the contrast in both PC3pip and PC3flu tumors was similar with both NB and PSMA-NB. At later time points PC3pip tumor show high contrast with PSMA-NB. (b) Mean time intensity curves (TIC) of PC3pip tumors, PC3flu tumors, and kidneys after IV bubble administration. The TIC data was collected from uniform regions of interest drawn on the acquired image stacks.

Both PC3pip and PC3flu tumors and kidneys were localized in the same field of view (FOV) using ultrasound B-mode imaging. Before bubble injection, tumors and kidneys are not visible under ultrasound in the nonlinear contrast harmonic imaging (CHI) mode (Fig. 3a). A total of 200 μl of undiluted NB (~8 × 10^10^ PSMA-NB or NB) were injected through the tail vein, and continuous contrast mode US was performed to visualize the bubble dynamic in the tumors and the kidney. Rapid enhancement of the contrast was observed first in the kidneys followed by both tumors approximately 30 sec to 2 min post-injection. The time to peak for PSMA-NB in the PC3pip tumor was slightly longer but it is not significantly different compared to that of PSMA-NB in PC3flu, (**Supplementary Fig. 3a**; 3.97 ± 1.20 min compared to 2.75 ± 1.31 min, P = 0.09). Furthermore, there was no significant difference between the time to peak for NB in either PC3pip or PC3flu tumor (P = 0.21). The average peak intensity was measured to be 15.64 ± 0.42 dB for PSMA-NB and NB in the PC3pip and PC3flu tumors and it is not significantly different in each case **(Supplementary Fig. 3b)**. The NB accumulation was compared and validated using clinically available MB-Lumason®. The peak intensities obtained for PSMA-NB (15.96 ± 1.83 dB) and NB (15.60 ± 2.53 dB) were significantly different from the one observed with the Lumason® (7.38 ± 0.33 dB, p < 0.001). Also, the duration of signal enhancement with Lumason® was limited to < 5 min indicating low stability and low circulation time of Lumason® MB in the blood stream.

Table 1 shows the quantitative parameters obtained from the time intensity curves, including peak enhancement, time to peak, area under the curve (AUC), and 25% of the maximum peak. Importantly, the AUC was calculated for wash in (WiAUC) and wash out (WoAUC) phases separately, since the main differences in bubble dynamics were expected during the washout phase. The total AUC is shown in **Supplementary Fig. 3c**. The WiAUC for all cases were consistent, however, the WoAUC of PSMA-NB in PC3pip tumor showed a significant, 2-fold increase, compared to all other groups (**Supplementary Fig. 3d**; p < 0.001). Since the TIC are similar in all cases at the early time points, but deviate at the later time points, the half peak maximum or the half time of peak intensity (t_50%_) was also similar between groups (data not shown). However, the time to reach 25% of the maximum peak intensity (t_75%_) for PSMA-NB in PC3pip tumor was 2-fold longer than all other groups (p < 0.01) (Table 1). These results indicated prolonged retention of targeted NB in PSMA-expressing tumors.

**Table 1.** Summary of kinetic parameters obtained from time intensity curve(TIC)

| Bubble Type | Tumor | Peak Time (min) | Peak Intensity (dB) | Time to Reach 25% of Peak – $t_{75\%}$(min) | Wash-in AUC | Washout AUC |
| --- | --- | --- | --- | --- | --- | --- |
| PSMA-NB | PC3pip | 3.95 ± 1.21 | 15.96 ± 1.83 | 20.17 ± 8.14* | 49.39 ± 14.54 | 200.84 ± 31.36* |
|  | PC3flu | 2.75 ± 1.31 | 15.60 ± 2.53 | 9.75 ± 3.76 | 40.32 ± 10.85 | 112.06 ± 30.47 |
| NB | PC3pip | 2.83 ± 1.90 | 15.94 ± 2.84 | 9.71 ± 3.91 | 39.53 ± 14.33 | 116.51 ± 25.26 |
|  | PC3flu | 2.66 ± 1.26 | 15.05 ± 2.55 | 8.33 ± 3.03 | 36.73 ± 13.09 | 102.17 ± 31.31 |

N=7, error bars represent mean ± s.d., \* P < 0.001.

To further clarify the effect that active nanobubble targeting on tumor accumulation, the signal from PSMA-NB was normalized to the signal from NB in both PC3pip and PC3flu tumors at selected time points. This was enabled by the fact that each mouse in the cohort received a randomized injection of both targeted and untargeted NBs. Thus, we can compare the effect of targeting, and of target expression in tissue, in addition to any unrelated confounding effects that may result from surface functionalization of the bubbles. Fig. 4a shows a significantly higher ratio (>2) at t = 15 min post-injection of bubbles in PC3pip tumor compared to the PC3flu tumor and was significantly different at each time point. The high ratio persisted, and in fact continued to increase, for up to 30 min. To minimize the effect of the variability in tumors between animal to animal, the signal from each bubble in both tumors was also normalized to the kidney signal of the same animal at each time points. The kidney represents “normal tissue” where no extravascular accumulation of the NBs would be expected. Fig. 4b again revealed consistently increasing signal only in the PSMA-NB in the PC3pip tumors. These bubbles showed >2-fold high normalized signal compared to that of all the other groups at the study endpoint.

**Figure 4.**
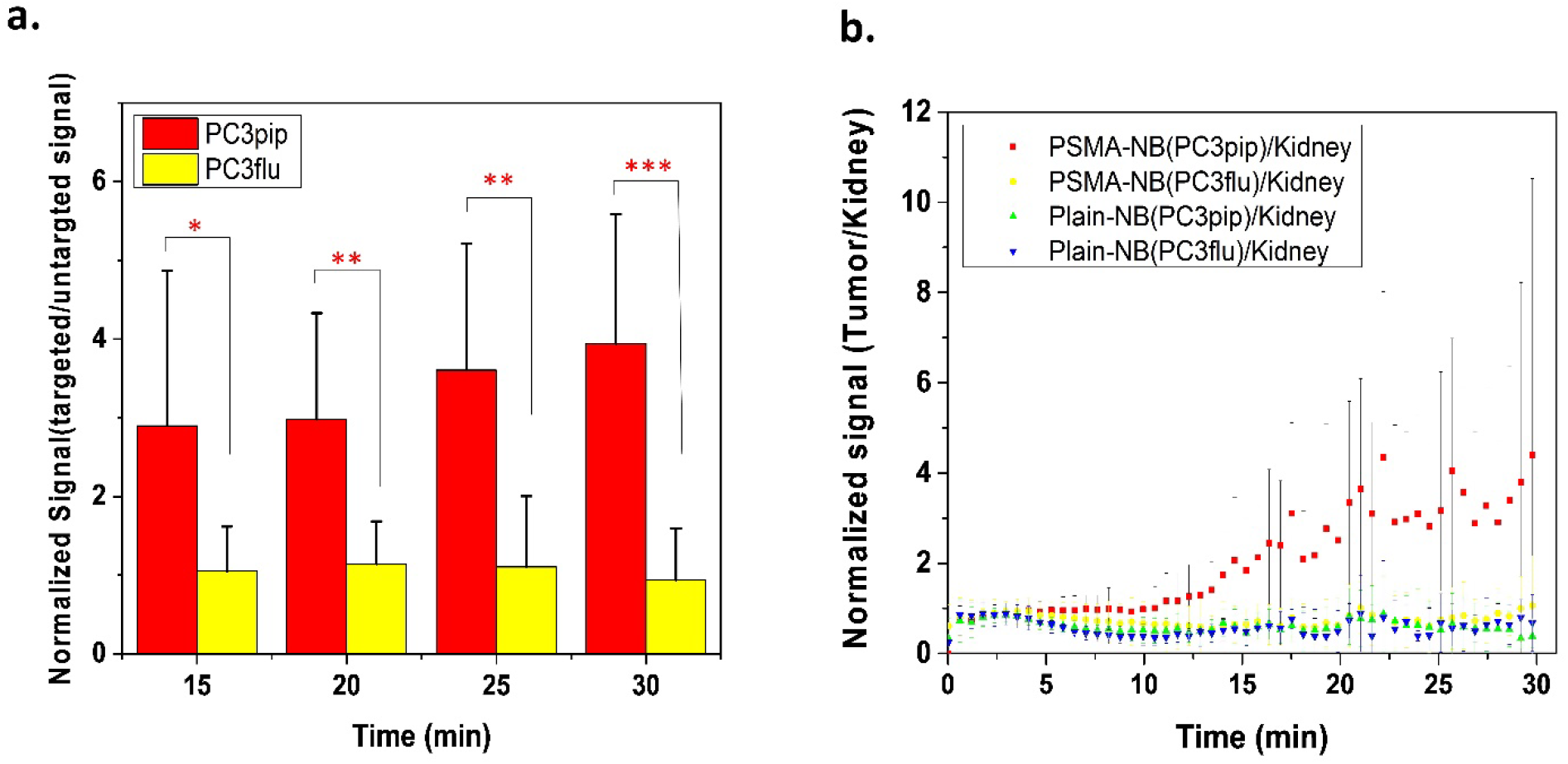
Normalized signal shows the high ratio with PSMA-NB in PC3pip tumor. (a) US signal obtained from PSMA-NB normalized to the signal from NB at each time point. (b) US signal obtained from PSMA-NB or NB normalized to the same bubble signal in kidney at each time point.

No significant differences were noted between PSMA-NB and NB groups for the time to peak, peak max, WiAUC in kidneys. Although the AUC in kidney with PSMA-NB injection was higher compared to the NB group, there is no significant difference between the two types of NBs (**Supplementary Fig. 3c)**. Moreover, in the kidneys there was no significant difference in t_50 %_ or t_75%_ values between PSMA-NB and NB (data not shown).

### Histology

To further validate that PSMA-targeted NB can extravasate into the tumor matrix, the bubbles were tagged with a fluorescent dye; Cy5.5, and injected via tail vein. Twenty-five min post-injection, the anesthetized animal was euthanized using cardiac perfusion with saline to remove all blood and concurrently all circulating bubbles. The tumors and kidneys were harvested for histological analysis. In animals harboring the PC3pip tumors, the Cy5.5-PSMA-NB signal can be seen deep in the tissue, distally from the microvasculature, providing strong evidence that the targeted NBs extravasate from the vasculature and enter the tumor interstitial space (Fig. 5a). The Cy5.5-PSMA-NB signal in PC3pip tumor was significantly higher (5.4 fold) compared to retained signal in tumors that did not express PSMA, PC3flu tumor (P < 0.001) (Fig.5b). Likewise, non-targeted Cy5.5-NB had low accumulation in both PC3pip and PC3flu tumors, which was comparable to targeted NB levels in the PC3flu tumors after perfusion. We noted no significant histological differences of vasculature between both types of tumors when sections were stained with anti-CD-31 (Fig 5b). Quantification of the Cy5.5 labeled bubbles revealed that the targeted bubbles showed 5-fold more accumulation and localization with the CD31 stained vasculature in the PC3pip compared to PC3flu tumors (Fig 5d). The overexpression of the PSMA in the PC3pip cells was confirmed by immunohistochemistry results (Fig. 5 c, e; 63.66± 1.51 vs 1.83± 0.15).

**Figure 5.**
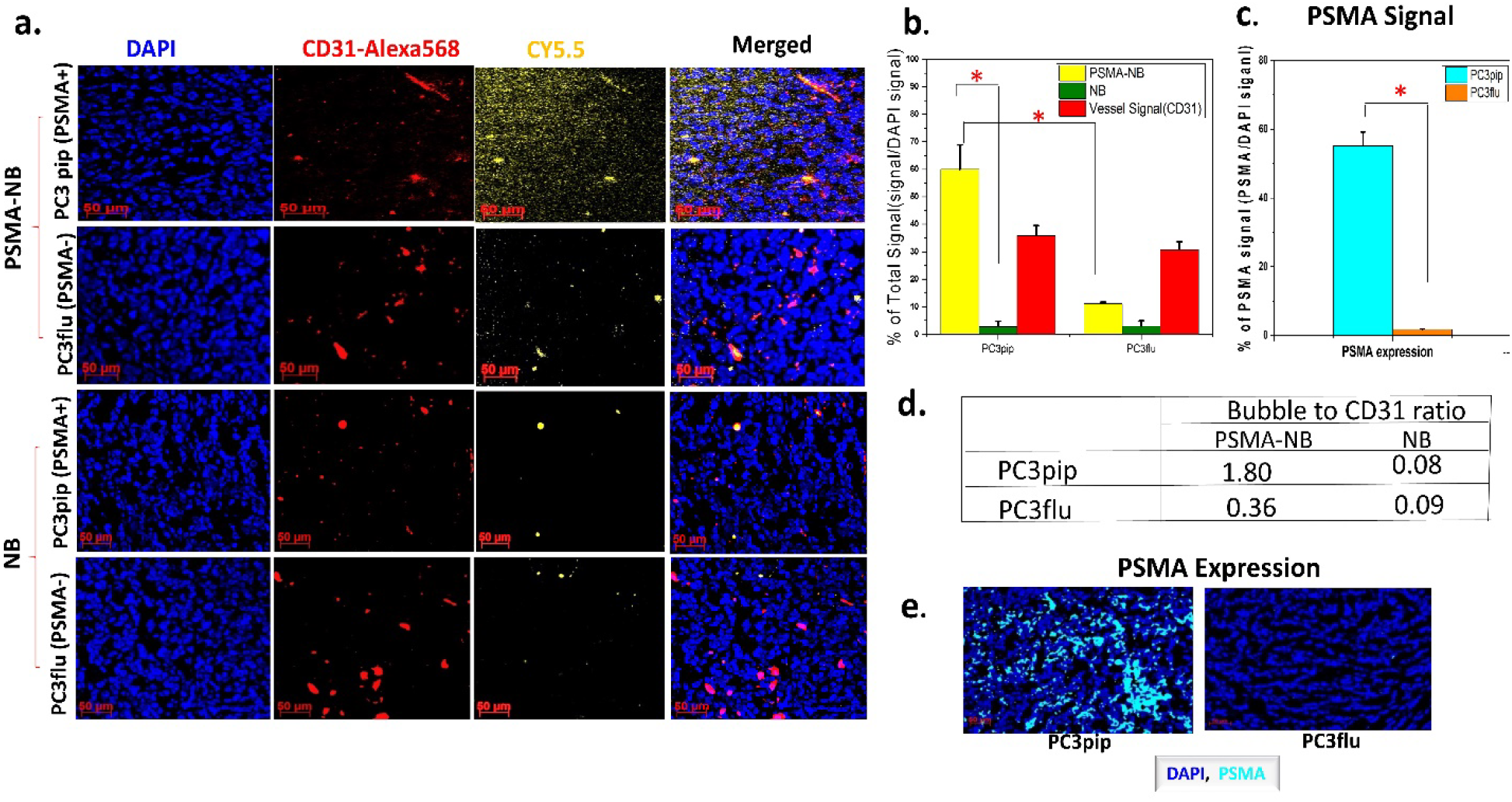
Histology images confirm the Cy5.5-PSMA-NB accumulation in PSMA-positive PC3pip tumor that were excised after cardiac perfusion with PBS. (a) Representative fluorescence images showing bubble distribution in tumor (orange). Cy5.5-PSMA-NB and showed higher extravasation compared to the Cy5.5-NB. Images show the extravasation of bubbles beyond the tumor vasculature (red). (b) The signal intensities of bubbles and vessel are shown here expressed as the percentage of total cell fluorescence in each tumor section. Cy5.5-PSMA-NB signal in PC3pip tumor is significantly higher from that of PC3flu tumor and the NB signal in PC3pip tumor. (c) The signal intensities of Cy5.5-PSMA-NB in both PC3pip and PC3flu tumors as the percentage of total cells of tumor tissues. Cy5.5-PSMA-NB signal in PC3pip tumor is significantly higher compared to that in PC3flu tumor. (d) The bubble to CD31 ratio in both PC3pip and PC3flu tumors. (e) Representative fluorescence images showing higher PSMA expression in PC3pip tumor compared to that in PC3flu tumor. N=3, error bars represent mean ± s.d., * P < 0.001.

## Discussion

Currently, PCa biopsies often provide false negative results. Consequently, there is a critical need for improved tools for PCa detection that can be used to better inform biopsy procedures. A wide range of imaging approaches is being examined to fill this unmet need, with MRI-guided biopsies and MRI preplanning showing the most promise^44–49^. Multiparametric MRI (mpMRI) and MRI-guided in gantry biopsies have been used in the clinic, and some sites have shown that the rate of PCa detection with these procedures is improved compared to transrectal US-guided biopsies^45–49^. However, MRI-based exams and procedures still present a number of challenges for broad adoption: they are not widely available, they are not portable and have a confined working space, and they require additional expertise outside of urology (e.g. radiology). The procedures can also be quite expensive and add significant time and cost to the diagnostic workflow. Finally, the negative predictive value (the ability to confidently state that a patient is malignancy-free), while hugely beneficial, has not been shown to be effective at this point with MRI ^50–51^.

A large body of work has instead focused on use of contrast enhanced US that are capable of molecular level imaging for this application. However, due to the rapid dissipation of encapsulated gas that causes a rapid signal decay, MB have a short *in vivo* half-life, which is not subtle enough to delineate pathological tissues from surrounding normal tissues for US-guided biopsies. Currently, the most promising MB tool is BR55, which targets VEGFR2^14–15, 52^, as discussed above, also has comparatively low specificity and sensitivity. While MBs are useful for visualizing vascular targets, they are restricted to the blood pool. In contrast, nano-sized particles may be able to penetrate leaky tumor vasculature to reach tissue targets located outside of vessels. In order to achieve sufficient dosage to gain a predominant signal at the target, the nanoparticles should outflow from the physical and biological barriers in the body, such as renal clearance, etc^53–55^. Formulation of NBs with a robust and resilient shell has prolonged their circulating time *in vivo*^24^. In the current study, in order to achieve specific US molecular imaging, the NBs were targeted to PSMA, which is expressed in high level in both androgen-dependent PCa and androgen-independent PCa, and a common target for PCa US imaging^9–10^. Consequently, PSMA-1 tagged NBs open an avenue for specific imaging at a molecular level in the tumor environment.

Bubble diameter and concentration obtained using resonant mass measurement indicated that both NB and the PSMA-NB are in nanoscale (< 300 nm) range and have a concentration several orders of magnitude higher than commercial MB agents. The small size and high shell deformability facilitated by inclusion of the edge activator, propylene glycol^56^ may enable NBs to pass through the neovasculature ^57–58^. The *in vitro* cell uptake studies provided the optimum concentration of PSMA-1 that can be incorporated into the NBs to achieve peak uptake. The optimum ligand density obtained (35 × 10^3^ ligands per NB) concurred with the reported values^43^. Cell uptake studies also demonstrated selectivity of binding of PSMA-NBs to the PSMA-positive PC3pip tumor cells.

In the *in vivo* acoustic evaluation, the nondestructive low mechanical index (MI = 0.1) was used to construct TIC and the parametric dynamic CEUS imaging was used to derive the contrast kinetics. Most importantly, this data demonstrates selective uptake of PSMA-NB in PSMA-positive PC3pip tumors compared to the PSMA-negative PC3flu tumors. In comparing the contribution of active targeting to this behavior it was crucial to also examine untargeted NB in the same tumors, thus mice received a randomized injection of these bubbles in the same exam, either before or after PSMA-NB administration. We observed that the time to peak and maximum peak enhancement were similar in both groups, signifying the similar dynamic of PSMA-NB and NB in the blood stream and also comparable morphology and vasculature in both PC3pip and PC3flu tumors. In stark contrast, the washout was slower with PSMA-NB in the PC3pip tumor compared to all other groups, as shown by persistent enhancement over 30 min, 2-fold higher WoAUC and t_75%_. Preferential accumulation of PSMA-NB in the PSMA expressing tumors was noticeable as early as 10 min after injection and increased over time. This is a considerable improvement over the relatively small and transient differences noted in prior studies of PSMA-targeted contrast agents^9–10^. Lumason® MB exhibited significant rapid enhancement in the kidney and relatively minor enhancement in tumors, and were cleared from the circulation rapidly without any significant binding. The high tumor to kidney signal ratio with PSMA-NB in PC3pip tumor also supports the conclusion that targeted NBs accumulate in PSMA-expressing tumors. However, to fully elucidate the fate of NB in the tumor tissue, a 3D imaging modality will be more informative compared to 2D imaging; these studies are currently ongoing.

Histological findings confirm the PSMA targeted NB can specifically recognize the tumors with PSMA expression. The percentage of PSMA-NB in PC3pip tumor is approximately 6-fold higher than that in PC3flu tumor. There bubbles were previously shown to also be intact and capable of generating acoustic activity, even following tumor perfusion^59^. The data also revealed that PSMA appears to be present in the tumor vasculature, which is a well-documented phenomenon for many tumors including some prostate cancers^60^. Concurrent with this we observed signal from PSMA-NB in the neovasculature in some PC3pip tumors even after the whole body perfusion of the animal (data not shown). Moreover, the histology data showed an increase of PSMA-NB accumulation in kidneys. While absolute levels were small, the PSMA-NB signal in the kidney was on average 6-fold higher compared to the NB signal (**Supplementary Fig. 4a, b)**. Notably, we observed positive signal from PSMA in kidney, yet approximately 30% lower compared to the PC3pip tumor (**Supplementary Fig. 4c).** Despite this unexpected expression, the kidney normalized data in Fig 4b above shows increasing enhancement in the PC3pip tumors over 30 min, indicating that the affinity, biding and retention of PSMA-NBs in tumors is considerable.

## Conclusion

The prolonged persistence of US signal of PSMA-NB in PCa provides exciting future opportunities to facilitate improved PCa detection with multiparametric contrast-enhanced ultrasound. The same principles can also make real-time PCa biopsies with US guidance a reality in the future. Additional studies are ongoing to also examine in detail extravasation of nanobubbles in tumors and quantify their acoustic activity following extravasation. Furthermore, in depth studies will elucidate the capability of PSMA-NB in delineating PCa using orthotropic PCa in mouse and larger animal models with the eventual goal of clinical translation into a rapid real-time biopsy guidance strategy.

## Supporting Information

Supporting information is available via the Web version of PubMed Central.

## Acknowledgements

This work was funded by the National Institutes of Health (1R01EB025741) and the Office of the Assistant Secretary of Defense for Health Affairs, through the Prostate Cancer Research Program under Award No. W81XWH-16-1-037I and W81XWH-16-1-0372. We also acknowledge additional support from the Case Comprehensive Cancer Center P30CA043703 in the form of a pilot grant and National Foundation for Cancer Research (NFCR). Views and opinions of, and endorsements by the author(s) do not reflect those of the National Institutes of Health or of the Department of Defense. We also thank Dr. C. Hernandez, A. Akhter, J. Lilly and Dr. B. Erokwu for their initial assistance with these studies.

## Methods

### Preparation of contrast agents

Lipid solution (10 mg/mL) for nanobubbles was prepared by first dissolving 1,2-dibehenoyl-snglycero-3-phosphocholine (C22, Avanti Polar Lipids Inc., Pelham, AL), 1,2 Dipalmitoyl-sn-Glycero-3-Phosphate (DPPA, Corden Pharma, Switzerland), 1,2-dipalmitoyl-sn-glycero-3-phosphoethanolamine (DPPE, Corden Pharma, Switzerland), and 1,2-distearoyl-snglycero-3-phosphoethanolamine-N-[methoxy(polyethylene glycol)-2000] (ammonium salt) (DSPE-mPEG 2000, Laysan Lipids, Arab, AL) with 6:1:2:1 ratio in propylene glycol (PG, Sigma Aldrich, Milwaukee, WI) by heating and sonicating at 80 °C until all the lipids were dissolved. Mixture of glycerol (Gly, Acros Organics) and phosphate buffer solution (0.8 mL, Gibco, pH 7.4) preheated to 80 °C was added to the lipid solution. The resulting solution was sonicated for 10 min at room temperature. Note that we typically prepared batches of 5 or 10 samples at a time and the amounts were adjusted as appropriate. For bubble formation, the solution (1 mL) was transferred to a 3 mL headspace vial, capped with a rubber septum and aluminum seal, and sealed with a vial crimper. Air was manually removed with a 30 mL syringe and was replaced by injecting octafluoropropane (C3F8, Electronic Fluorocarbons, LLC, PA) gas. After air was replaced by C3F8, the phospholipid solution was activated by mechanical shaking with a VialMix shaker (Bristol-Myers Squibb Medical Imaging Inc., N. Billerica, MA) for 45s. Nanobubbles were isolated from the mixture of foam and microbubbles by centrifugation at 50 rcf for 5 mins with the headspace vial inverted, and the 100 μL NB solution withdrawn from a fixed distance of 5 mm from the bottom with a 21G needle^24^.

PSMA-NB were prepared by adding DSPE-PEG-PSMA-1 (25μg/ml) to the initial lipid solution and followed the above protocol. To prepare DSPE-PEG-PSMA-1, PSMA-1 (from prof. James Basilion lab) was mixed with DSPE-PEG-MAL (1,2-distearoyl-snglycero-3-phosphoethanolamine-N-[methoxy(polyethylene glycol)-2000-Maleimide, Laysan Bio, Arab, AL) in 1:2 ratio at pH 8.0 in PBS. After combined, the mixture was vortexed thoroughly and was reacted for 4 hours on the vial rotator at 4 °C. The product was lyophilized and the resultant powder was dissolved in PBS to obtain DSPE-PEG-PSMA-1 stock solution.

### Size, concentration, and surface charge of NBs and non-buoyant particles

The size distribution and concentration of NBs were characterized with resonant mass measurement (Archimedes®, Malvern Panalytical) equipped with a nanosensor capable of measuring particle size between 50 nm and 2000 nm. The nanosensor was pre-calibrated with 565 nm polystyrene beads (Thermo ScientificTM Nanospehere Size Standards 3560A). The NB solution was diluted with PBS (500×) to obtain an acceptable limit of detection (< 0.01 Hz) and coincidence (< 5%). Prior to any measurement, a 5-min PBS blank was run to ensure that the system fluidics and sensor were free of particles. Also, between measurements the sensor and microfluidic tubing were rinsed for 30 seconds with PBS followed by 2 “sneezes” for at least 3 cycles. During the sample measurement, NB solution was loaded for 120 seconds and analyzed at 2 and 5 psi, respectively. Samples measurement was finalized after 1000 particles were measured. Data was exported from the Archimedes software (version 1.2) and analyzed for positive and negative counts^24^. Dilution was accounted for in calculating the NB concentration. Surface charge of the diluted NB solution (500X) was measure with an Anton Paar LitesizerTM 500.

### Cell Culture

Retrovirally transformed PSMA positive PC3pip cells and transfection control PC3flu cells were originally obtained from Dr. Michel Sadelain in the year 2000 (Laboratory of Gene Transfer and Gene Expression, Gene Transfer and Somatic Cell Engineering Facility, Memorial-Sloan Kettering Cancer Center, New York, NY). The two cell lines were last checked and authenticated by western blot in 2017. Cells were grown at 37°C and 5% CO_2_ under a humidified atmosphere. Cells were maintained in RPMI 1640 medium supplemented (Invitrogen Life Technology, Grand Island, NY) with 2 mM L-glutamine and 10% Fetal Bovine Serum.

### Celluar Uptake Studies

PC3pip and PC3flu cells were plated on coverslips at about 70% confluency. Rhodamine labeled NBs were prepared by mixing DSPE-Rhodamine (50μl) into the lipid cocktail solution that used to make NB. Twenty-four hours later, cells were incubated with Rhodamine B-labeled nanobubbles for 1 hour. After incubation, cells were washed three times with PBS, fixed with 4% paraformaldehyde, counterstained with 2-(4-amidinophenyl)-6-indolecarbamidine dihydrochloride (DAPI), mounted with Fluor-Mount aqueous mounting solution, and observed under Leica DM4000B fluorescence microscopy (Leica Microsystem Inc, Buffalo Grove, IL). The fluorescent intensity was then quantified by Image J.

### Animal models

Animals were handled according to a protocol approved by the Institutional Animal Care and Use Committee (IACUC) at Case Western Reserve University and were in accordance with all applicable protocols and guidelines in regards to animal use. Four to six week old male athymic nude mice were anesthetized with inhalation of 3% isoflurane with 1L/min oxygen and were implanted subcutaneously with 1×10^6^ of PSMA-negative PC3flu and PSMA-positive PC3pip cells in 100 µL matrigel. Animals were observed every other day until tumors reached at about 8-10 mm in diameter.

### Pharmacokinetic study

Two weeks after inoculation, the tumor diameter reached 1 cm animals were used in the study (n=7). The US probe (PLT-1204BT, AplioXG SSA-790A, Toshiba Medical Imaging Systems, Otawara-Shi, Japan) was placed to visualize the ultrasound images of the PC3pip and PC3flu dual tumors and the kidney in the same field of view. Two hundred μl of either undiluted Plain NB or PSMA-NB NB were administrated via tail vein. After injection of NB, the change of tissue contrast was measured using contrast harmonic imaging (CHI, frequency 12.0MHz; MI, 0.1; dynamic range, 65dB; gain, 70dB; imaging frame rate, 0.2 frames/s). The images were acquired in raw data format as a function of time. Fifteen seconds after raw data acquisition started, nanobubble solution was administrated and continuous image acquisition continued for 30 min. The remaining NBs were burst by repeated flash replenish (high energy pulses). Thirty minutes later (1h after first injection, the contrast was reached to baseline level) the same mouse received PSMA-NB or plain-NB respectively (n=7). Lumason® (sulfur hexafluoride lipid-type A microspheres, Bracco Diagnostics Inc.) were tested *in vivo* (n=3). Lumason® was prepared according to the protocol provided by the manufacturer. The raw data were processed with software provided by the scanner manufacturer. The kidney, and tumor areas were delineated by drawing regions of interest (ROIs) and the signal intensity in each ROI as a function of time (time-intensity curve - TIC) was calculated. The data were exported to Excel, the baseline was subtracted from TIC, and the calculated peak value of TIC was used to normalize the data to obtain the decay of signal.

### Histological Analysis

Animals were divided into 3 groups: Cy5.5-PSMA-NB (n =3), Cy5.5-NB (n = 3), and no contrast control. Cy 5.5 labeled NBs were prepared by mixing DSPE-PEG-Cy 5.5 (100μl) into the lipid cocktail solution that used to make NB. Mice received either 200 μl of undiluted contrast material or PBS alone via tail vein. Twenty five minutes after contrast agent injection, animals were scan using US to detect the US signal and then PBS perfusion was performed with 50ml PBS though left ventricle. After perfusion tumors were scan again to perceive the US signal that generate from intact NB. Both PSMA (+) and PSMA (-) tumors and the kidney were harvested, fixed in paraformaldehyde and embedded in optimal cutting temperature compound (OCT Sakura Finetek USA Inc., Torrance, CA). The tissues were cut into 8 um slices. Then the CD31 staining was performed to visualize the tumor vessels. Briefly, tissues were wash 3 times with PBS and incubate with protein blocking solution that contain 0.5% Triton X-100 (Fisher Scientific, Hampton, NH). Then tissues were incubated in 1:250 diluted primary antibody (CD31 (PECAM-1) Monoclonal Antibody (390) Fisher Scientific, Hampton, NH) for 24 hrs at 4° C. After washed with PBS, one-hour incubation of Alexa 568 tagged secondary antibody (Fisher Scientific, Hampton, NH) performed and stained with DAPI (Vecor Laboratories, Burlingame, CA) using standard techniques. The fluorescence images were obtained and analyzed using Axio Vision V 4.8.1.0, Carl Zeiss software (Thornwood, NY). For PSMA immunohistochemistry, tissues were wash 3 times with PBS and incubate with protein blocking solution (Thermo Fisher Scientific, Waltham, MA). Then tissues were incubated in 1:150 diluted PSMA primary antibody (Thermo Fisher Scientific, Waltham, MA) for 24 hrs at 4 ° C and followed the above steps as for CD31 staining.

### Statistical Analysis

Graphs and statistical analyses were generated using Microsoft Excel and Origin lab. Unpaired Student’s t-test (two-tailed) was used to compare two groups. Data are presented as a mean ± STD (standard deviation). The experiments were repeated at least three times for each experiment, unless stated otherwise.

## References

1. Negoita, S.; Feuer, E. J.; Mariotto, A.; Cronin, K. A.; Petkov, V. I.; Hussey, S. K.; Benard, V.; Henley, S. J.; Anderson, R. N.; Fedewa, S.; Sherman, R. L.; Kohler, B. A.; Dearmon, B. J.; Lake, A. J.; Ma, J.; Richardson, L. C.; Jemal, A.; Penberthy, L., Annual Report to the Nation on the Status of Cancer, part II: Recent changes in prostate cancer trends and disease characteristics. Cancer 2018, 124 (13), 2801–2814.

2. Cronin, K. A.; Lake, A. J.; Scott, S.; Sherman, R. L.; Noone, A. M.; Howlader, N.; Henley, S. J.; Anderson, R. N.; Firth, A. U.; Ma, J.; Kohler, B. A.; Jemal, A., Annual Report to the Nation on the Status of Cancer, part I: National cancer statistics. Cancer 2018, 124 (13), 2785–2800.

3. Mottet, N.; Bellmunt, J.; Bolla, M.; Briers, E.; Cumberbatch, M. G.; De Santis, M.; Fossati, N.; Gross, T.; Henry, A. M.; Joniau, S.; Lam, T. B.; Mason, M. D.; Matveev, V. B.; Moldovan, P. C.; van den Bergh, R. C. N.; Van den Broeck, T.; van der Poel, H. G.; van der Kwast, T. H.; Rouviere, O.; Schoots, I. G.; Wiegel, T.; Cornford, P., EAU-ESTRO-SIOG Guidelines on Prostate Cancer. Part 1: Screening, Diagnosis, and Local Treatment with Curative Intent. Eur Urol 2017, 71 (4), 618–629.

4. Roethke, M.; Anastasiadis, A. G.; Lichy, M.; Werner, M.; Wagner, P.; Kruck, S.; Claussen, C. D.; Stenzl, A.; Schlemmer, H. P.; Schilling, D., MRI-guided prostate biopsy detects clinically significant cancer: analysis of a cohort of 100 patients after previous negative TRUS biopsy. World J Urol 2012, 30 (2), 213–8.

5. Pallwein, L.; Mitterberger, M.; Pelzer, A.; Bartsch, G.; Strasser, H.; Pinggera, G. M.; Aigner, F.; Gradl, J.; Zur Nedden, D.; Frauscher, F., Ultrasound of prostate cancer: recent advances. Eur Radiol 2008, 18 (4), 707–15.

6. Smeenge, M.; Barentsz, J.; Cosgrove, D.; de la Rosette, J.; de Reijke, T.; Eggener, S.; Frauscher, F.; Kovacs, G.; Matin, S. F.; Mischi, M.; Pinto, P.; Rastinehad, A.; Rouviere, O.; Salomon, G.; Polascik, T.; Walz, J.; Wijkstra, H.; Marberger, M., Role of transrectal ultrasonography (TRUS) in focal therapy of prostate cancer: report from a Consensus Panel. BJU Int 2012, 110 (7), 942–8.

7. Smeenge, M.; de la Rosette, J. J.; Wijkstra, H., Current status of transrectal ultrasound techniques in prostate cancer. Curr Opin Urol 2012, 22 (4), 297–302.

8. Sanna, V.; Pintus, G.; Bandiera, P.; Anedda, R.; Punzoni, S.; Sanna, B.; Migaleddu, V.; Uzzau, S.; Sechi, M., Development of polymeric microbubbles targeted to prostate-specific membrane antigen as prototype of novel ultrasound contrast agents. Mol Pharm 2011, 8 (3), 748–57.

9. Wang, L.; Li, L.; Guo, Y.; Tong, H.; Fan, X.; Ding, J.; Huang, H., Construction and in vitro/in vivo targeting of PSMA-targeted nanoscale microbubbles in prostate cancer. Prostate 2013, 73 (11), 1147–58.

10. Fan, X.; Wang, L.; Guo, Y.; Tu, Z.; Li, L.; Tong, H.; Xu, Y.; Li, R.; Fang, K., Ultrasonic Nanobubbles Carrying Anti-PSMA Nanobody: Construction and Application in Prostate Cancer-Targeted Imaging. PLoS One 2015, 10 (6), e0127419.

11. Zlitni, A.; Yin, M.; Janzen, N.; Chatterjee, S.; Lisok, A.; Gabrielson, K. L.; Nimmagadda, S.; Pomper, M. G.; Foster, F. S.; Valliant, J. F., Development of prostate specific membrane antigen targeted ultrasound microbubbles using bioorthogonal chemistry. PLoS One 2017, 12 (5), e0176958.

12. Hernot, S.; Unnikrishnan, S.; Du, Z.; Shevchenko, T.; Cosyns, B.; Broisat, A.; Toczek, J.; Caveliers, V.; Muyldermans, S.; Lahoutte, T.; Klibanov, A. L.; Devoogdt, N., Nanobody-coupled microbubbles as novel molecular tracer. J Control Release 2012, 158 (2), 346–53.

13. Willmann, J. K.; Bonomo, L.; Carla Testa, A.; Rinaldi, P.; Rindi, G.; Valluru, K. S.; Petrone, G.; Martini, M.; Lutz, A. M.; Gambhir, S. S., Ultrasound Molecular Imaging With BR55 in Patients With Breast and Ovarian Lesions: First-in-Human Results. J Clin Oncol 2017, 35 (19), 2133–2140.

14. Tardy, I.; Pochon, S.; Theraulaz, M.; Emmel, P.; Passantino, L.; Tranquart, F.; Schneider, M., Ultrasound molecular imaging of VEGFR2 in a rat prostate tumor model using BR55. Invest Radiol 2010, 45 (10), 573–8.

15. Pochon, S.; Tardy, I.; Bussat, P.; Bettinger, T.; Brochot, J.; von Wronski, M.; Passantino, L.; Schneider, M., BR55: a lipopeptide-based VEGFR2-targeted ultrasound contrast agent for molecular imaging of angiogenesis. Invest Radiol 2010, 45 (2), 89–95.

16. Smeenge, M.; Tranquart, F.; Mannaerts, C. K.; de Reijke, T. M.; van de Vijver, M. J.; Laguna, M. P.; Pochon, S.; de la Rosette, J.; Wijkstra, H., First-in-Human Ultrasound Molecular Imaging With a VEGFR2-Specific Ultrasound Molecular Contrast Agent (BR55) in Prostate Cancer: A Safety and Feasibility Pilot Study. Invest Radiol 2017, 52 (7), 419–427.

17. Baur, A. D. J.; Schwabe, J.; Rogasch, J.; Maxeiner, A.; Penzkofer, T.; Stephan, C.; Rudl, M.; Hamm, B.; Jung, E. M.; Fischer, T., A direct comparison of contrast-enhanced ultrasound and dynamic contrast-enhanced magnetic resonance imaging for prostate cancer detection and prediction of aggressiveness. Eur Radiol 2018, 28 (5), 1949–1960.

18. de Leon, A.; Perera, R.; Nittayacharn, P.; Cooley, M.; Jung, O.; Exner, A. A., Ultrasound Contrast Agents and Delivery Systems in Cancer Detection and Therapy. Adv Cancer Res 2018, 139, 57–84.

19. Gao, Y.; Hernandez, C.; Yuan, H. X.; Lilly, J.; Kota, P.; Zhou, H.; Wu, H.; Exner, A. A., Ultrasound molecular imaging of ovarian cancer with CA-125 targeted nanobubble contrast agents. Nanomedicine 2017, 13 (7), 2159–2168.

20. Perera, R. H.; Wu, H.; Peiris, P.; Hernandez, C.; Burke, A.; Zhang, H.; Exner, A. A., Improving performance of nanoscale ultrasound contrast agents using N,N-diethylacrylamide stabilization. Nanomedicine 2017, 13 (1), 59–67.

21. Perera, R. H.; Hernandez, C.; Zhou, H.; Kota, P.; Burke, A.; Exner, A. A., Ultrasound imaging beyond the vasculature with new generation contrast agents. Wiley Interdiscip Rev Nanomed Nanobiotechnol 2015, 7 (4), 593–608.

22. Perera, R. H.; Solorio, L.; Wu, H.; Gangolli, M.; Silverman, E.; Hernandez, C.; Peiris, P. M.; Broome, A. M.; Exner, A. A., Nanobubble ultrasound contrast agents for enhanced delivery of thermal sensitizer to tumors undergoing radiofrequency ablation. Pharmaceutical research 2014, 31 (6), 1407–17.

23. Wu, H.; Rognin, N. G.; Krupka, T. M.; Solorio, L.; Yoshiara, H.; Guenette, G.; Sanders, C.; Kamiyama, N.; Exner, A. A., Acoustic characterization and pharmacokinetic analyses of new nanobubble ultrasound contrast agents. Ultrasound in medicine & biology 2013, 39 (11), 2137–46.

24. Hernandez, C.; Abenojar, E. C.; Hadley, J.; Leon, A. C. d.; Coyne, R.; Perera, R.; Gopalakrishnan, R.; Basilion, J. P.; Kolios, M. C.; Exner, A. A., Sink or float? Characterization of shell-stabilized bulk nanobubbles using a resonant mass measurement technique. Nanoscale 2018, Under press.

25. Maeda, H.; Nakamura, H.; Fang, J., The EPR effect for macromolecular drug delivery to solid tumors: Improvement of tumor uptake, lowering of systemic toxicity, and distinct tumor imaging in vivo. Advanced drug delivery reviews 2013, 65 (1), 71–9.

26. Shi, J.; Kantoff, P. W.; Wooster, R.; Farokhzad, O. C., Cancer nanomedicine: progress, challenges and opportunities. Nat Rev Cancer 2017, 17 (1), 20–37.

27. Kobayashi, H.; Watanabe, R.; Choyke, P. L., Improving conventional enhanced permeability and retention (EPR) effects; what is the appropriate target? Theranostics 2013, 4 (1), 81–9.

28. Castanares, M. A.; Mukherjee, A.; Chowdhury, W. H.; Liu, M.; Chen, Y.; Mease, R. C.; Wang, Y.; Rodriguez, R.; Lupold, S. E.; Pomper, M. G., Evaluation of prostate-specific membrane antigen as an imaging reporter. J Nucl Med 2014, 55 (5), 805–11.

29. Genady, A. R.; Janzen, N.; Banevicius, L.; El-Gamal, M.; El-Zaria, M. E.; Valliant, J. F., Preparation and Evaluation of Radiolabeled Antibody Recruiting Small Molecules That Target Prostate-Specific Membrane Antigen for Combined Radiotherapy and Immunotherapy. J Med Chem 2016, 59 (6), 2660–73.

30. Barrett, J. A.; Coleman, R. E.; Goldsmith, S. J.; Vallabhajosula, S.; Petry, N. A.; Cho, S.; Armor, T.; Stubbs, J. B.; Maresca, K. P.; Stabin, M. G.; Joyal, J. L.; Eckelman, W. C.; Babich, J. W., First-in-man evaluation of 2 high-affinity PSMA-avid small molecules for imaging prostate cancer. J Nucl Med 2013, 54 (3), 380–7.

31. Ghosh, A.; Heston, W. D., Tumor target prostate specific membrane antigen (PSMA) and its regulation in prostate cancer. J Cell Biochem 2004, 91 (3), 528–39.

32. Sweat, S. D.; Pacelli, A.; Murphy, G. P.; Bostwick, D. G., Prostate-specific membrane antigen expression is greatest in prostate adenocarcinoma and lymph node metastases. Urology 1998, 52 (4), 637–40.

33. Ristau, B. T.; O’Keefe, D. S.; Bacich, D. J., The prostate-specific membrane antigen: lessons and current clinical implications from 20 years of research. Urol Oncol 2014, 32 (3), 272–9.

34. Perner, S.; Hofer, M. D.; Kim, R.; Shah, R. B.; Li, H.; Moller, P.; Hautmann, R. E.; Gschwend, J. E.; Kuefer, R.; Rubin, M. A., Prostate-specific membrane antigen expression as a predictor of prostate cancer progression. Hum Pathol 2007, 38 (5), 696–701.

35. Kawakami, M.; Okaneya, T.; Furihata, K.; Nishizawa, O.; Katsuyama, T., Detection of prostate cancer cells circulating in peripheral blood by reverse transcription-PCR for hKLK2. Cancer Res 1997, 57 (19), 4167–70.

36. Mhawech-Fauceglia, P.; Zhang, S.; Terracciano, L.; Sauter, G.; Chadhuri, A.; Herrmann, F. R.; Penetrante, R., Prostate-specific membrane antigen (PSMA) protein expression in normal and neoplastic tissues and its sensitivity and specificity in prostate adenocarcinoma: an immunohistochemical study using mutiple tumour tissue microarray technique. Histopathology 2007, 50 (4), 472–83.

37. Wang, X.; Huang, S. S.; Heston, W. D.; Guo, H.; Wang, B. C.; Basilion, J. P., Development of targeted near-infrared imaging agents for prostate cancer. Mol Cancer Ther 2014, 13 (11), 2595–606.

38. Wang, X.; Tsui, B.; Ramamurthy, G.; Zhang, P.; Meyers, J.; Kenney, M. E.; Kiechle, J.; Ponsky, L.; Basilion, J. P., Theranostic Agents for Photodynamic Therapy of Prostate Cancer by Targeting Prostate-Specific Membrane Antigen. Mol Cancer Ther 2016, 15 (8), 1834–44.

39. Lee, J.; Shen, W.; Payer, K.; Burg, T. P.; Manalis, S. R., Toward attogram mass measurements in solution with suspended nanochannel resonators. Nano Lett 2010, 10 (7), 2537–42.

40. Burg, T. P.; Godin, M.; Knudsen, S. M.; Shen, W.; Carlson, G.; Foster, J. S.; Babcock, K.; Manalis, S. R., Weighing of biomolecules, single cells and single nanoparticles in fluid. Nature 2007, 446 (7139), 1066–9.

41. Olcum, S.; Cermak, N.; Wasserman, S. C.; Christine, K. S.; Atsumi, H.; Payer, K. R.; Shen, W.; Lee, J.; Belcher, A. M.; Bhatia, S. N.; Manalis, S. R., Weighing nanoparticles in solution at the attogram scale. Proc Natl Acad Sci U S A 2014, 111 (4), 1310–5.

42. Segers, T.; Lohse, D.; Versluis, M.; Frinking, P., Universal Equations for the Coalescence Probability and Long-Term Size Stability of Phospholipid-Coated Monodisperse Microbubbles Formed by Flow Focusing. Langmuir 2017, 33 (39), 10329–10339.

43. Elias, D. R.; Poloukhtine, A.; Popik, V.; Tsourkas, A., Effect of ligand density, receptor density, and nanoparticle size on cell targeting. Nanomedicine 2013, 9 (2), 194–201.

44. Smeenge, M.; Tranquart, F.; Mannaerts, C. K.; de Reijke, T. M.; van de Vijver, M. J.; Laguna, M. P.; Pochon, S.; de la Rosette, J. J.; Wijkstra, H., First-in-human ultrasound molecular imaging with a VEGFR2-specific ultrasound molecular contrast agent (BR55) in prostate cancer: a safety and feasibility pilot study. Investigative radiology 2017, 52 (7), 419–427.

45. Schouten, M. G.; Hoeks, C. M.; Bomers, J. G.; Hulsbergen-van de Kaa, C. A.; Witjes, J. A.; Thompson, L. C.; Rovers, M. M.; Barentsz, J. O.; Fütterer, J. J., Location of Prostate Cancers Determined by Multiparametric and MRI-Guided Biopsy in Patients With Elevated Prostate-Specific Antigen Level and at Least One Negative Transrectal Ultrasound–Guided Biopsy. American Journal of Roentgenology 2015, 205 (1), 57–63.

46. Schoots, I. G.; Roobol, M. J.; Nieboer, D.; Bangma, C. H.; Steyerberg, E. W.; Hunink, M. M., Magnetic resonance imaging–targeted biopsy may enhance the diagnostic accuracy of significant prostate cancer detection compared to standard transrectal ultrasound-guided biopsy: a systematic review and meta-analysis. European urology 2015, 68 (3), 438–450.

47. Arsov, C.; Rabenalt, R.; Blondin, D.; Quentin, M.; Hiester, A.; Godehardt, E.; Gabbert, H. E.; Becker, N.; Antoch, G.; Albers, P., Prospective randomized trial comparing magnetic resonance imaging (MRI)-guided in-bore biopsy to MRI-ultrasound fusion and transrectal ultrasound-guided prostate biopsy in patients with prior negative biopsies. European urology 2015, 68 (4), 713–720.

48. Filson, C. P.; Natarajan, S.; Margolis, D. J.; Huang, J.; Lieu, P.; Dorey, F. J.; Reiter, R. E.; Marks, L. S., Prostate cancer detection with magnetic resonance-ultrasound fusion biopsy: The role of systematic and targeted biopsies. Cancer 2016, 122 (6), 884–892.

49. Siddiqui, M. M.; Rais-Bahrami, S.; Turkbey, B.; George, A. K.; Rothwax, J.; Shakir, N.; Okoro, C.; Raskolnikov, D.; Parnes, H. L.; Linehan, W. M., Comparison of MR/ultrasound fusion–guided biopsy with ultrasound-guided biopsy for the diagnosis of prostate cancer. Jama 2015, 313 (4), 390–397.

50. Kaufmann, S.; Kruck, S.; Kramer, U.; Gatidis, S.; Stenzl, A.; Roethke, M.; Scharpf, M.; Schilling, D., Direct comparison of targeted MRI-guided biopsy with systematic transrectal ultrasound-guided biopsy in patients with previous negative prostate biopsies. Urol Int 2015, 94 (3), 319–25.

51. Afshar-Oromieh, A.; Haberkorn, U.; Schlemmer, H. P.; Fenchel, M.; Eder, M.; Eisenhut, M.; Hadaschik, B. A.; Kopp-Schneider, A.; Rothke, M., Comparison of PET/CT and PET/MRI hybrid systems using a 68Ga-labelled PSMA ligand for the diagnosis of recurrent prostate cancer: initial experience. Eur J Nucl Med Mol Imaging 2014, 41 (5), 887–97.

52. Abou-Elkacem, L.; Bachawal, S. V.; Willmann, J. K., Ultrasound molecular imaging: Moving toward clinical translation. Eur J Radiol 2015, 84 (9), 1685–93.

53. Nichols, J. W.; Bae, Y. H., Odyssey of a cancer nanoparticle: from injection site to site of action. Nano Today 2012, 7 (6), 606–618.

54. Florence, A. T., “Targeting” nanoparticles: the constraints of physical laws and physical barriers. J Control Release 2012, 164 (2), 115–24.

55. Yu, M.; Zheng, J., Clearance Pathways and Tumor Targeting of Imaging Nanoparticles. ACS Nano 2015, 9 (7), 6655–74.

56. Lee, E. H.; Kim, A.; Oh, Y. K.; Kim, C. K., Effect of edge activators on the formation and transfection efficiency of ultradeformable liposomes. Biomaterials 2005, 26 (2), 205–10.

57. Maeda, H.; Bharate, G. Y.; Daruwalla, J., Polymeric drugs for efficient tumor-targeted drug delivery based on EPR-effect. Eur J Pharm Biopharm 2009, 71 (3), 409–19.

58. Hobbs, S. K.; Monsky, W. L.; Yuan, F.; Roberts, W. G.; Griffith, L.; Torchilin, V. P.; Jain, R. K., Regulation of transport pathways in tumor vessels: role of tumor type and microenvironment. Proc Natl Acad Sci U S A 1998, 95 (8), 4607–12.

59. Perera, R.; De Leon, A. C.; Wang, X.; Ramamurthy, G.; Peiris, P.; Basilion, J.; Exner, A. A., Nanobubble Extravasation in Prostate Tumors Imaged with Ultrasound: Role of Active versus Passive Targeting. EEE Int. Ultrason. Symp. IUS 2018, In press.

60. Chang, S. S.; Reuter, V. E.; Heston, W. D.; Bander, N. H.; Grauer, L. S.; Gaudin, P. B., Five different anti-prostate-specific membrane antigen (PSMA) antibodies confirm PSMA expression in tumor-associated neovasculature. Cancer Res 1999, 59 (13), 3192–8.

