## Supporting Information- Supplemental Figures for "Real Time Ultrasound Molecular Imaging of Prostate Cancer with PSMA-targeted Nanobubbles"

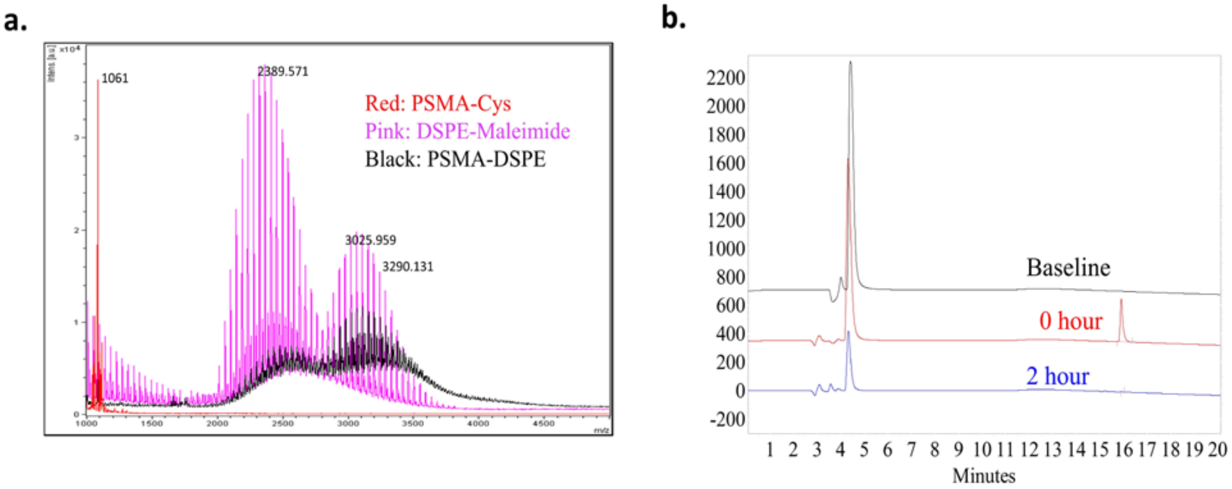


**Figure S1**. (a) MALDI-TOF MS analysis of DSPE-PSMA conjugation (b) HPLC analysis of PSMA-1 and DSPE-PSMA conjugation.

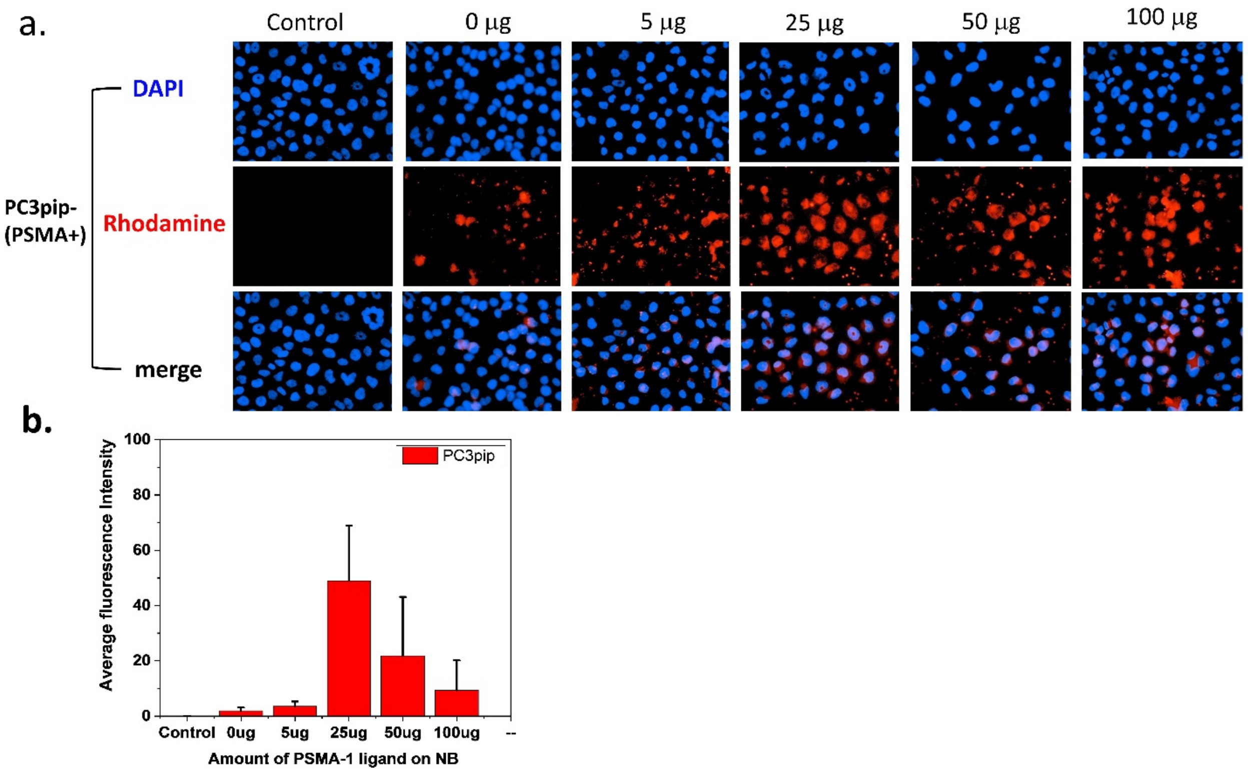


**Figure S2**. *In vitro* cellular uptake experiments reveal PSMA-NB selectively bind to the PSMA - positive PC3pip cells. (a) PSMA-positive PC3pip cells on coverslips were incubated with no NB (control), Rhodamine-NB or Rhodamine-PSMA-NB with different PSMA amounts for 1hr. Nuclei were stained using DAPI (blue) and uptake of Rhodamine tagged NB and PSMA-NB (red) was assessed by fluorescence microscopy. Images were taken at 40X. Representative images are shown from three independent experiments. (b) Optimization of PSMA amount in NB. Twenty-five µg of PSMA-1 shows significantly higher fluorescence signal. n=3, error bars represent mean ± s.d., * $P < 0.05$

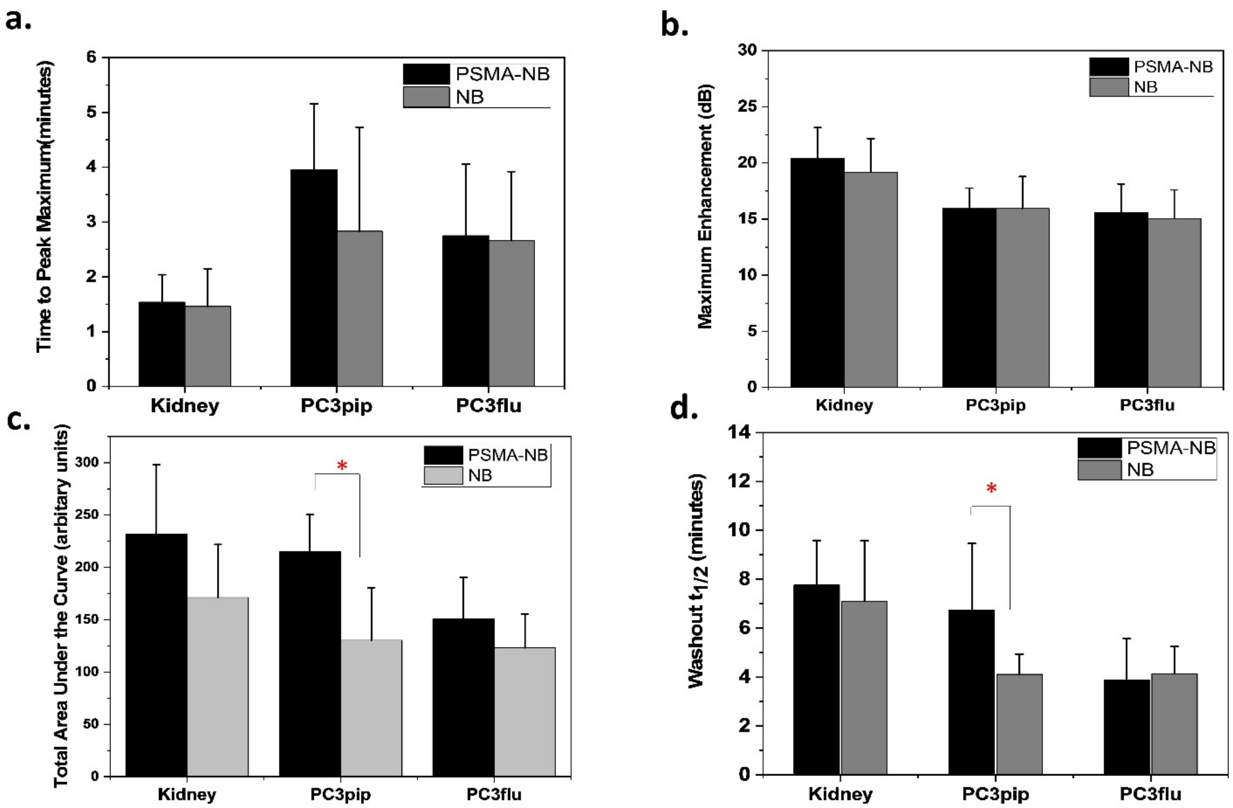


**Figure S3.** Quantitative *in vivo* parameters obtained from TIC analysis comparing kinetics of targeted (PSMA-NB) and untargeted (NB) bubbles. (a) Time to peak enhancement, (b) Maximum enhancement, (c) Total area under the curve (AUC), (d) Washout half-life. Significant differences ($p<0.05$) were seen in the AUC and washout half-life between targeted and untargeted bubbles in the PSMA-expressing tumors (PC3pip).

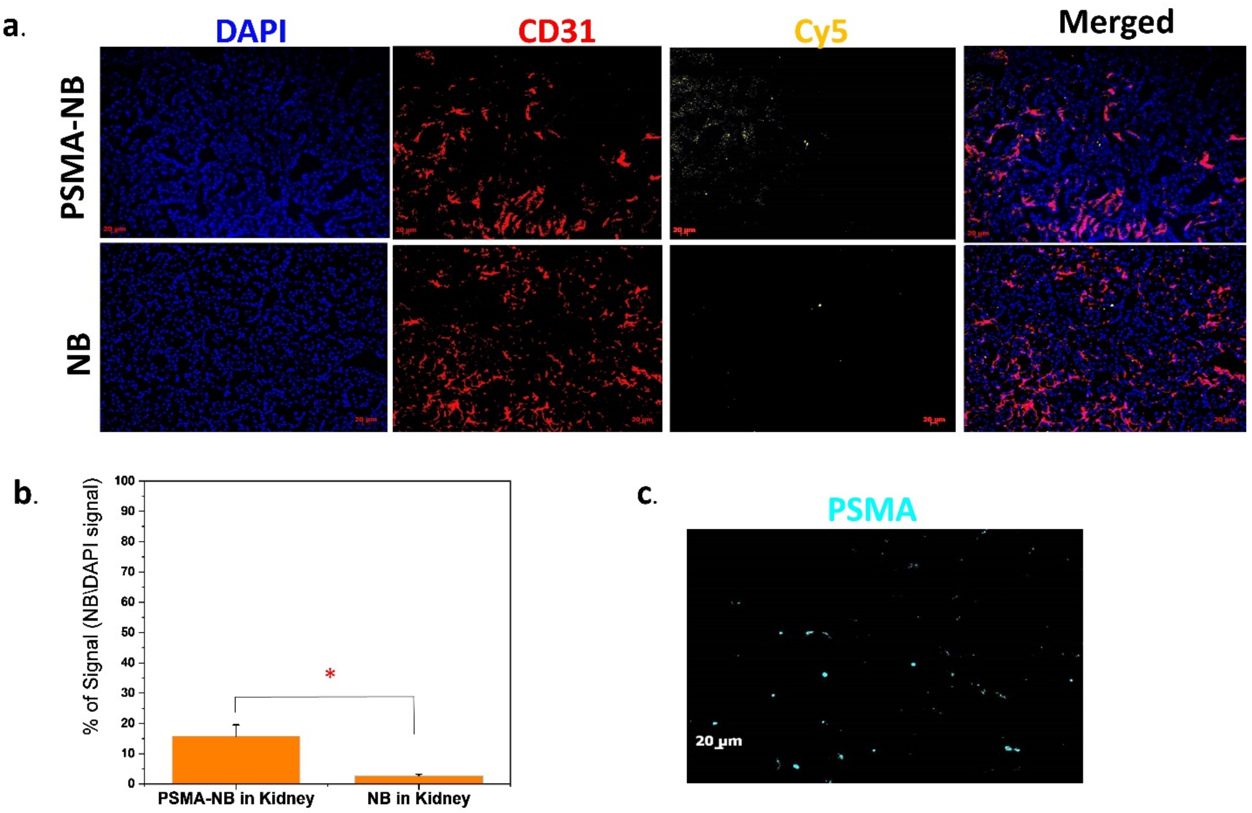


**Figure S4.** (a) Representative fluorescence images showing the bubble (yellow) and vasculature (red) distribution in Kidney. Residual Cy5.5 tagged PSMA-NB signal were seen in the kidney and its presence appears to be correlated to the location of vasculature (red). (b) The Cy5.5 signal intensity was quantified in these images and is shown here as a percentage of total cells in each kidney section field of view. PSMA-NB Cy5.5 signal was significantly different from the NB signal. (c) Representative fluorescence images showing the PSMA expression in the kidney (cyan).
